# Fuzzy RNA-recognition by the *Trypanosoma brucei* editosome

**DOI:** 10.1101/2021.06.10.446919

**Authors:** Wolf-Matthias Leeder, H. Ulrich Göringer

## Abstract

The recognition of RNA-molecules by proteins and protein complexes is a critical step on all levels of gene expression. Typically, the generated ribonucleoprotein complexes rely on the binary interaction of defined RNA-sequences or precisely folded RNA-motifs with dedicated RNA-binding domains on the protein side. Here we describe a new molecular recognition principle of RNA-molecules by a high molecular mass protein complex. By chemically probing the solvent accessibility of mitochondrial pre-mRNAs when bound to the *Trypanosoma brucei* editosome we identified multiple similar but nonidentical RNA-motifs as editosome contact sites. However, by treating the different motifs as mathematical graph objects we demonstrate that they fit a consensus 2D-graph consisting of 4 vertices (*V*) and 3 edges (*E*) with a Laplacian eigenvalue of 0.523 (λ_2_). We establish that a synthetic 4*V*(3*E*)-RNA is sufficient to compete for the editosomal pre-mRNA binding site and that it is able to inhibit RNA-editing *in vitro*. Our analysis corroborates that the editosome has adapted to the structural multiplicity of the mitochondrial mRNA-folding space by recognizing a fuzzy continuum of RNA-folds that fit a consensus graph-descriptor. This provides a mechanism on how the protein complex is able to bind the structurally pleomorphic pool of pre- and partially edited mRNAs. We speculate that other fuzzy RNA-recognition motifs exist especially for proteins that interact with multiple RNA-species.

## Introduction

Mitochondrial gene expression in the protozoan parasite *Trypanosoma brucei* relies on a nucleotide-specific RNA-editing reaction. In the process sequence deficient and as a consequence non-functional primary transcripts are re-modeled to functional messenger (m)RNAs by the site-specific insertion and deletion of uridine (U)-nucleotides (nt) (reviewed in Cruz-Reyes et al., 2018). Depending on the transcript the extend of the reaction can vary drastically. While only 4 U-nt are inserted into the cytochrome oxidase II (COII) transcript, more than 600 U-nt are processed in the case of the NADH-dehydrogenase subunit 7 (ND7) mRNA. For nine, so-called pan-edited transcripts more than 50% of the mature mRNA-sequence are the result of the processing reaction and since edited mRNAs code for key components of the mitochondrial ribosome and the mitochondrial electron transport and chemiosmosis system the RNA-processing reaction is essential for the organism.

Catalytic machinery of the process is a mitochondrial multiprotein complex termed the editosome. The 800kDa particle provides a catalytic surface for all steps of the reaction cycle including endo- and exonuclease, terminal uridylyl-transferase (TUTase), RNA-ligase and RNA-chaperone activities (reviewed in Göringer, 2012). Accessory enzymes contribute as well and involve RNA-annealing (Müller et al., 2001; Müller and Göringer, 2002, Mehta et al., 2020) and RNA-helicase activities (Missel et al., 1997; Li et al., 2011, Kumar et al., 2020). The sequence specificity for the U-insertion/U-deletion reaction is provided by 40-60nt long, non-coding (nc) RNAs termed guide (g)RNAs (Blum et al., 1990). Guide RNAs hybridize to the pre- and partially edited mRNAs and control the U-nt insertion/deletion reaction by antiparallel base pairing. At steady state more than 1000 different gRNAs are expressed in the mitochondria of insect-stage African trypanosomes (Koslowsky et al., 2014) and in the majority of cases multiple (≤10) gRNAs are needed to fully edit a single pre-mRNA.

Initial step of the editing reaction is the binding of a pre-edited transcript into the single RNA-binding site of the editosome (Böhm et al., 2012). The reaction is assisted by multiple oligonucleotide/oligosaccharide-binding (OB)-fold proteins on the surface of the editosome, which collectively execute a chaperone-type RNA-unfolding activity (Voigt et al., 2018). The activity is driven by the intrinsically disordered protein (IDP)-domains of the different polypeptides (Czerwoniec et al., 2015) and as a result, the highly folded pre-mRNAs (Leeder et al., 2016a) become partially unfolded thereby increasing their probability to interact with a gRNA-molecule to initiate the reaction cycle (Voigt et al., 2018, Leeder et al., 2106b). Pre-mRNA/editosome complexes can be formed *in vitro* (Böhm et al., 2012) and pre-mRNA mimicking oligoribonucleotides become accurately edited in a gRNA-dependent manner (Del Campo et al., 2020). Together, this demonstrates that the pre-mRNA binding reaction is an editosome-inherent trait, which, at least *in vitro*, can be executed in the absence of additional protein factors. Intriguingly, the editosomal RNA-binding capacity is characterized by a staggering plasticity. For instance, editosomes process a highly pleomorphic set of RNA-ligands. This includes pre-edited mRNAs varying in length from 164nt (CR3) to 1117nt (Cyb) and literally thousands of partially edited mRNAs (Zimmer et al., 2018). Short, synthetic oligoribonucleotides of only 30nt are edited as well, as are phosphorothioate- and/or 2’-CH_3_O-modified RNA-oligonucleotides (Del Campo et al., 2020). Even more perplexing, editosomes maintain their RNA-binding capacity despite the fact that the pre-mRNAs continuously change their primary sequence by being edited. Pan-edited transcripts double their U-content from 30% to 60% during the processing reaction thereby altering their R/Y-ratio from R-rich (R/Y=1.9) to Y-rich (R/Y=0.6). How editosomes solve this “moving-target-problem” is not understood (Supplementary Fig. 1). In the same way, the RNA-binding motif that facilitates binding to the editosome is not known.

As demonstrated for many RNA/protein complexes, complex formation often results in the burial of parts of the RNA-surface deep inside the protein contact site (Jankowsky and Harris, 2015; Balcerak et al., 2019). As a consequence, these RNA-sequences become secluded from the surrounding solvent and can be identified by comparing the RNA-solvent accessibility in the free and protein-bound state. Here we determine the changes in solvent accessibility of five pre-edited mRNAs when bound to the *T. brucei* editosome using hydroxyl radical footprinting (HRP) (Fig. 1A and Supplementary Fig. 2). Hydroxyl radicals cleave the sugar-phosphate backbone of accessible ribonucleotides and with a size of only 0.36nm they are perfectly suited for solvent probing experiments (Costa and Monachello, 2014; Balasubramanian et al., 1998). We performed high-throughput HRP-experiments by identifying HR-cleavage positions as abortive cDNA-synthesis products using fluorophore-derivatized oligodesoxynucleotide primer molecules in combination with capillary electrophoresis (CE) and laser-induced fluorescense (LIF) detection (Duncan and Weeks, 2010a,b) (Fig. 1A). We identify one or two editosome-dependent RNA-footprints in all five pre-mRNAs. The different signatures are similar but not identical, which suggests a malleable, fuzzy-type RNA-recognition mode by the editosome similar to what has been described for fuzzy protein/protein complexes (Sharma et al., 2015). Using graph-theory (Schlick, 2018) we characterize the fuzzy recognition motif to consist of 4 vertices (*V*) and 3 edges (*E*) and confirm that a synthetic 4*V*(3*E*)-RNA is sufficient to compete for the editosomal pre-mRNA binding site and that it is able to inhibit RNA-editing *in vitro*. Our analysis demonstrates that the *T. brucei* editosome is competent to process the structural multiplicity or dynamic disorder in the pre- and partially edited mRNA-folding space by recognizing a fuzzy continuum of RNA-binding motifs. This rationalizes the enigmatic RNA-binding characteristics of the *T. brucei* editosome and provides a simple mechanism on how the high molecular mass protein complex is able to bind and process the structurally pleomorphic pool of pre- and partially edited mitochondrial mRNAs.

**Fig. 1.**
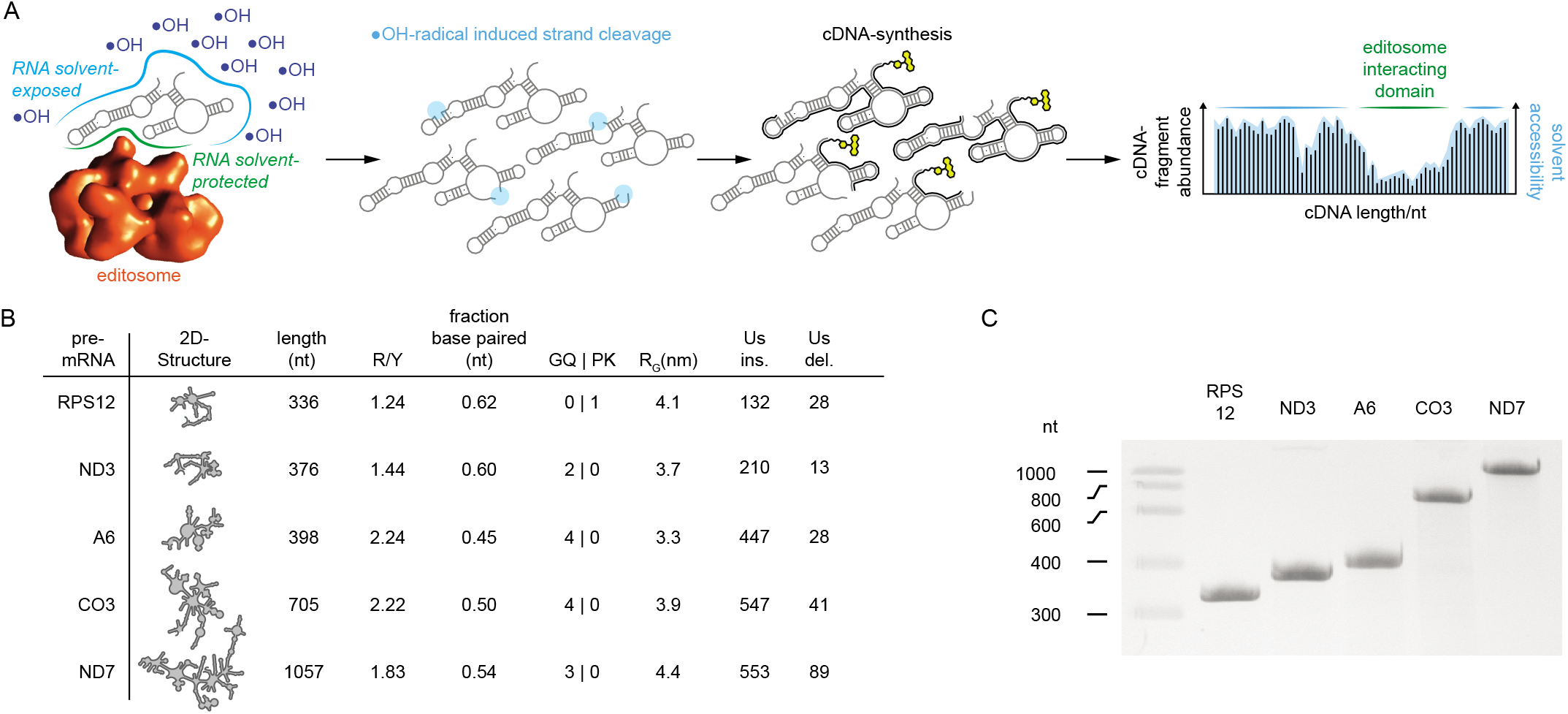
Hydroxyl radical footprinting of editosome/pre-mRNA-complexes. (A) 2D-representation of a mitochondrial pre-mRNA bound to the *T. brucei* editosome (red sphere). Complex formation generates an RNA/protein interface (green line), in which parts of the RNA become solvent-protected. Treatment of the complexes with hydroxyl radicals (•OH) results in RNA-backbone cleavage in all solvent-exposed RNA-domains (cyan line). At single-hit conditions, RNA-molecules are cleaved only once (circles in light blue) and strand cleavage positions are mapped by abortive cDNA-synthesis (black lines) using fluorophore-labeled primer molecules (yellow). cDNA-fragments are separated by capillary electrophoresis and quantified by laser-induced fluorescence. A plot of the cDNA-fragment abundance in relation to the nt-length represents a read-out of the nt-solvent accessibility. (B) Characteristics of the 5 tested *T. brucei* pre-mRNAs including 2D-structure, nt-length, purine/pyrimidine (R/Y)-ratio, fraction of base paired nucleotides, presence of G-quadruplex (GQ)- and pseudoknot (PK)-folds, radius of gyration (R_G_) and the number inserted (ins) and deleted (del) U-residues. (C) Electrophoretic separation of *in vitro* transcribed *T. brucei* pre-mRNAs RPS12, ND3, A6, CO3 and ND7. RNA-preparations (3μg) are separated in a 6% (w/v), urea-containing (8M) polyacrylamide gel followed by Toluidine Blue O staining. Samples purities are ≥97%.

## Results

### Probing the solvent accessibility of *T. brucei* pre-mRNAs when bound to the editosome

Of the 12 edited transcripts in the mitochondria of *T. brucei* we selected five pre-mRNAs for the analysis: the pre-mRNA of subunit 6 of the mitochondrial ATPase (A6), the transcript encoding subunit 3 of the cytochrome c oxidase (CO3), the pre-mRNAs of subunits 3 and 7 of the NADH dehydrogenase (ND3, ND7) and the transcript of ribosomal protein S12 (RPS12). All five RNAs represent extensively edited transcripts and cover a size range from 336nt (RPS12) to 1057nt (ND7) (Fig. 1B). The different RNAs were synthesized by run-off *in vitro* transcription with purities ≥98% (Fig. 1C). Following synthesis, the RNAs were refolded and editosome: pre-mRNA complexes were formed at a 2-fold molar excess of editosomes to ascertain that every RNA-molecule is editosome-bound. The RNA-solvent accessibility was analyzed by hydroxyl radical probing (HRP) (Duncan and Weeks, 2010a,b). Reactions were performed at single-hit conditions and sites of HR-induced RNA-backbone cleavage were identified as abortive cDNA-synthesis products. Cleavage intensities were normalized to a scale from 0 to 4 with a value of 1.0 representing the median cleavage intensity. Values ≤0.5 represent solvent inaccessible ribonucleotides (Duncan and Weeks, 2010a,b).

*In toto* we probed 5130 nt-positions comparing the five pre-mRNAs in their free and editosome-bound states. As shown in Fig. 2 all pre-mRNAs display complex solvent accessibility profiles both, as free RNAs and when bound to the editosome. The different HRP-traces are characterized by alternating regions of cleaved backbone positions above and below the median. When filtered with a moving average over a 3nt-window, these sequences span between 4-5nt in the free RNAs and 6nt in the editosome-bound states. Importantly, the HRP-profiles of the two folding states (free RNA *vs*. editosome-bound RNA) differ for every pre-mRNA. This is reflected in a mean Pearson correlation coefficient of *p*=0.75 between the two folding situations, which indicates that the solvent-accessibility of the 5 pre-mRNAs changes upon editosome binding. Localized differences can reach up to 3-times the median reactivity. To further analyze the data, we generated HRP-difference (ΔHRP)-plots by subtracting the nt-cleavage intensities of the free RNAs from the corresponding values of the editosome-bound state. This identifies solvent inaccessible nt-positions as negative values and nt-positions with increased solvent accessibility as positive values. Fig. 2 shows the ΔHRP-plots for all 5 pre-mRNAs. In every case solvent-protected nucleotides as well as nucleotide positions with enhanced solvent-accessibility can be identified and are dispersed over the entire primary sequences. This is consistent with the documented complex-inherent RNA-chaperone activity of the editosome (Böhm et al., 2012; Leeder et al., 2016b), which catalyzes a partial RNA-unfolding reaction to assist the formation of pre-mRNA/gRNA hybrid molecules to initiate the editing process (Voigt et al., 2018). Furthermore, by plotting the number of highly protected nucleotides in the editosome-bound state as a function of all interrogated nucleotides in the different RNAs we identified a saturation-type of behavior (Supplementary Fig. 3). A number of about 60nt are maximally involved in the pre-mRNA: editosome interaction, which suggests that the RNA-contact site of the editosome is a finite surface area.

**Fig. 2.**
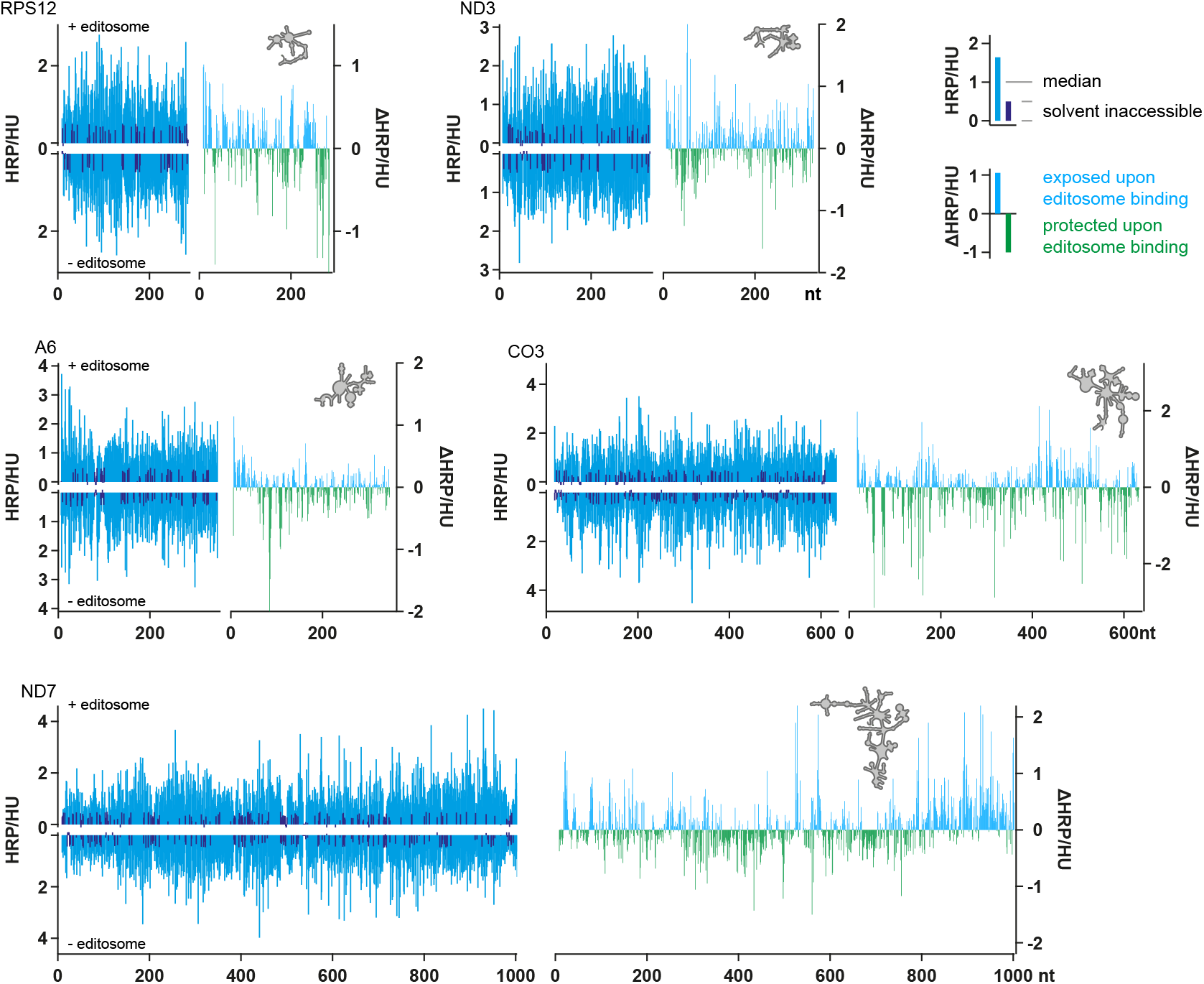
HRP-reactivity profiles. Left panels: Histograms of the mapped HRP-cleavage intensities (blue bars) *versus* nucleotide position for the RPS12-, ND3-, CO3-, A6- and ND7-pre-mRNAs in the presence (top) and absence (bottom) of editosomes. The data are normalized to the median reactivity and intensities ≤0.5 are considered solvent inaccessible. Right panels: Difference (Δ)HRP-plots of the cleavage intensities in the editosome-bound state minus cleavage intensities in the free RNA. Nucleotide positions that become protected upon editosome binding are in green and nucleotides with enhanced HRP-reactivities in blue. The 2D-structures of the different pre-mRNAs are shown as grey silhouettes. Replicates of the HRP-experiments correlate with Pearson correlation coefficients (*p*) ≥0.9. HU=HRP-unit.

### Fuzzy RNA-structure recognition by the editosome

Fig. 3 shows the experimentally-derived 2D minimal-free-energy (MFE)-structures of all tested pre-mRNAs. Plotted onto the different structures are the individual ΔHRP-values with nucleotide positions that become protected from HR-cleavage upon editosome binding in green and nucleotides with increased solvent accessibility upon complex formation shaded in blue. The data are suggestive of a clustering of all affected nucleotides on the 2Dstructure level especially for the majority of solvent inaccessible nucleotides. For the two large pre-mRNAs ND7 (1057nt) and CO3 (705nt) two protected 2D-RNA elements can be distinguished. For the 336nt RPS12-transcript and the A6- (398nt) and ND3 pre-mRNAs (376nt) single motifs are discernable. The individual RNA-domains have a length between 40-60nt. They differ in their base pairing pattern and are characterized by a mean thermodynamic stability (ΔG) of −0.45 kcal/mol/nt and a configurational entropy of 0.55. Almost all of them branch off from multi-loop arrangements and consist of irregular hairpins that contain next to multiple (38%) G:U bp asymmetric bulges and bulged-out nucleotides. All attempts to identify a shared primary or secondary structure in the protected sequences failed. No individual nucleotide, nor a specific nucleotide sequence or 2D-motif is over- or under-represented in any of the footprinted sequences (Fig. 4) despite the fact that the footprints are highly reproducible. Multiple experiments with different pre-mRNA preparations (≥5) and different editosome isolates (≥5) compare with a mean Pearson correlation coefficient (*p*) of 0.96 proving that the identified footprints are not stochastically driven. Instead, they seem to indicate a high degree of conformational plasticity or in other words “fuzziness”.

**Fig. 3.**
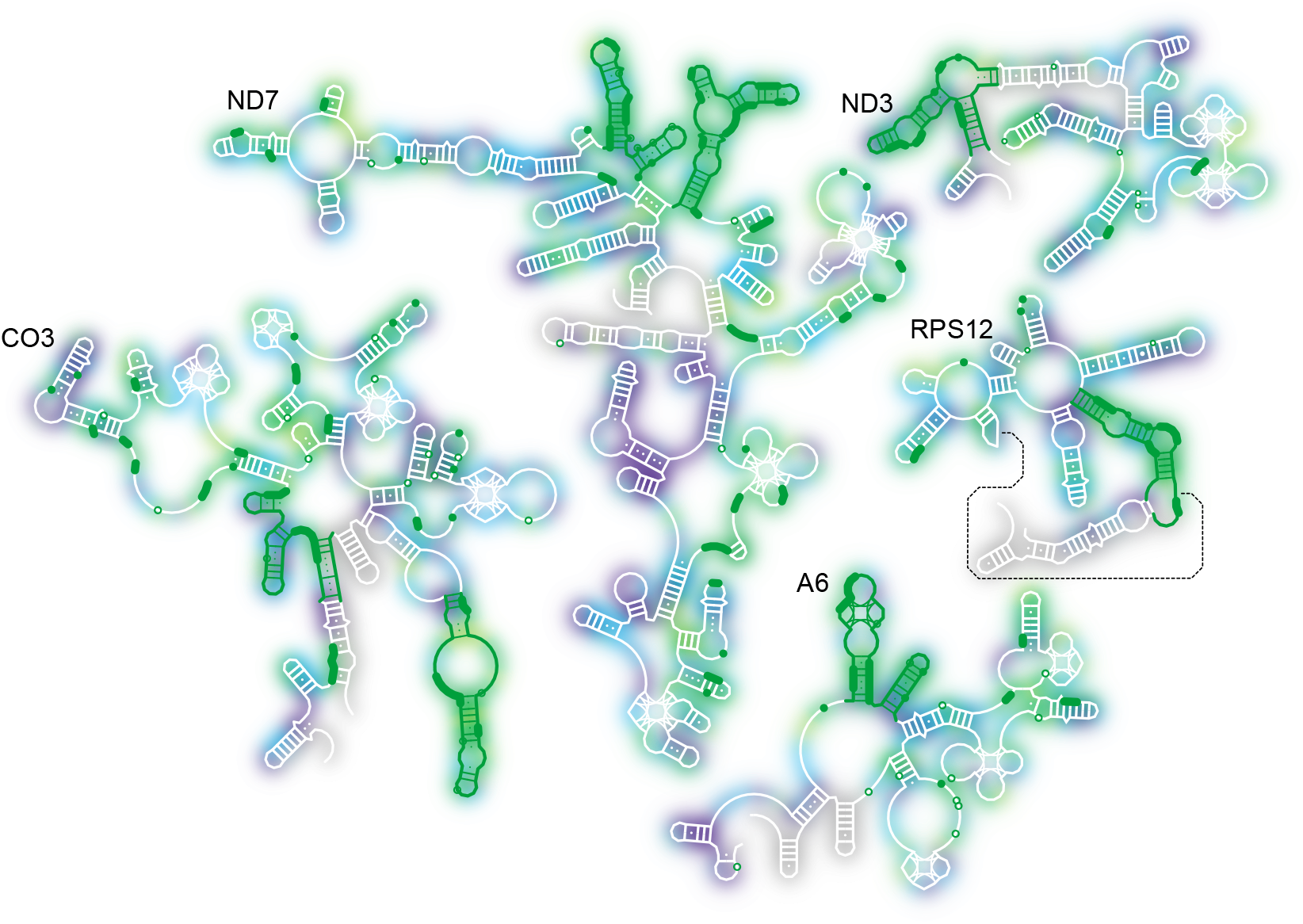
Secondary structure mapping of HRP-reactivities. Experimentally derived 2D-structures of the *T. brucei* pre-mRNAs encoding RPS12, ND3, A6, CO3 and ND7 (Leeder et al., 2016a). Non-canonical and GU-bp are shown as white dots. G-quadruplex elements are represented as leaf-like structures. A pseudoknot interaction in the RPS12 pre-mRNA is shown as a dotted line. Nucleotides protected from HRP-cleavage upon editosome binding (25^th^-percentile of negative ΔHRP-values x3) are marked in green and nucleotides with increased solvent accessibility upon editosome binding (75^th^-percentile of positive ΔHRP-values x3) are shaded/blurred in blue.

**Fig. 4.**
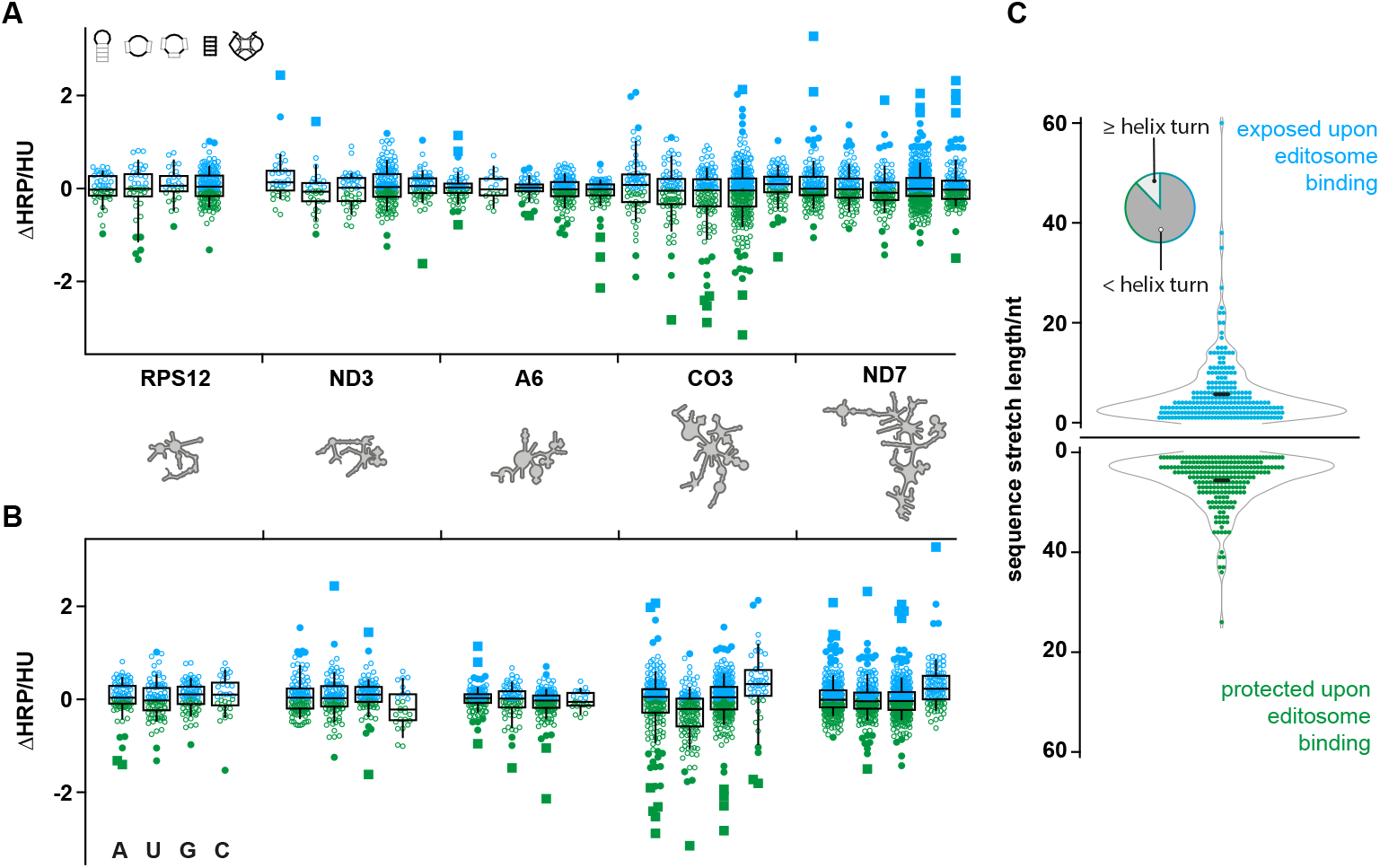
Statistical analysis of HRP-reactivity profiles. (A) Boxplot analysis of the ΔHRP-data for nucleotides in hairpin-loops, internal-loops, multi-loops, helices and GQ-elements (from left to right) for all tested pre-mRNAs (RPS12, ND3, A6, CO3, ND7). Sketches of the different 2D-elements are depicted in the in upper left corner. Blue: increased solvent accessibility upon editosome binding. Green: decreased solvent accessibility (protection) upon editosome binding. Whiskers indicate the 9^th^ and 91^st^ percentile. Outliers are shown as filled circles (1.5x the interquartile range (IQR)) or as filled squares (3x the IQR). The median is represented by a horizontal line. HU=HRP-unit. (B) The same plot as in (A) analysing the nucleotide identity (A, U, G, C from left to right). (C) Length distribution of continuous sequence stretches with increased (blue) or decreased (green) solvent accessibility upon editosome binding after smoothing over a 3nt window. Solvent-exposed and solvent-protected sequence stretches are on average 6nt long (black horizontal bar), which is equal to half a helical turn. Only 13% of the sequence stretches are ≥11nt (upper left pie chart). Further statistical data are provided in Supplementary Fig. 3 and Fig. 4.

### Graph theory-based RNA-motif analysis

To quantitatively assess the topological characteristics of the different editosome-interacting RNA-motifs we scrutinized the protected RNA-sequences in a coarse-grained analysis. For that we used a graph theory-based approach as established by Schlick and colleagues (Gan et al., 2003; Gan et al., 2004; Izzo et al., 2011; Schlick, 2018). The method transforms the identified RNA 2D-elements (Fig. 5A) into discrete mathematical graph representations (RNA-graphs) by describing the 2D-motifs as a set of vertices (*V*=loops, bulges, junctions) and edges (*E*=helices) (Fig. 5B). This shrinks the complexity of the identified RNA-elements down to their skeletal connectivity features and enables a quantitative comparison of the different RNA-elements using spectral graph theory-methods (Cvetkovic et al., 1995; Gan et al., 2004). The matrix of a graph specifies the degree of connectivity between the individual vertices. As suggested by Gan et al., 2004 we used the Laplacian *V* x *V*-matrix (*L*), which was constructed from the adjacency (*A*) and square diagonal (*D*) matrices of the graphs as *L*=*D-A. A* specifies the number of edges that connect pairs of vertices and *D* details the connectivity of each vertex (Fig. 5C). Furthermore, a *V*-vertex graph is characterized by its eigenvalue spectrum (0≤ λ_1_≤ λ_2_≤ λ_3_≤ λ_4_...). This can be used to calculate graph similarities. Typically, RNA-graphs are characterized by λ_1_=0 and λ_2_>0 (Gan et al., 2004). Thus, λ_2_ reveals the overall connectivity of a graph and graphs with similar λ_2_-values have similar topologies (Fig. 5C). Lastly, we concatenated the individual adjacency (*A*)- and square diagonal (*D*)-matrices to generate 3-dimensional *A*- and *D*-matrix stacks. The stacks were averaged along the 3^rd^-dimension to calculate mean *A*- and *D*-matrices (Fig. 5D). The resulting averaged *L*-matrix embodies a 4*V*(3*E*)-graph with an eigenvalue of 0.5228 (λ_2_). Examples of two 4*V*(3*E*)-tree graphs, after weighting the individual vertices, are shown in Fig. 5D and a typical outline of a 4*V*(3*E*) RNA 2D-fold is shown in Fig. 5E. The RNA has a three-way-junction topology in which three, mostly Watson-Crick paired helical elements, are linked by single-stranded RNA-sequences (Lescoute and Westhof, 2006; De la Peña et al., 2009).

**Fig. 5.**
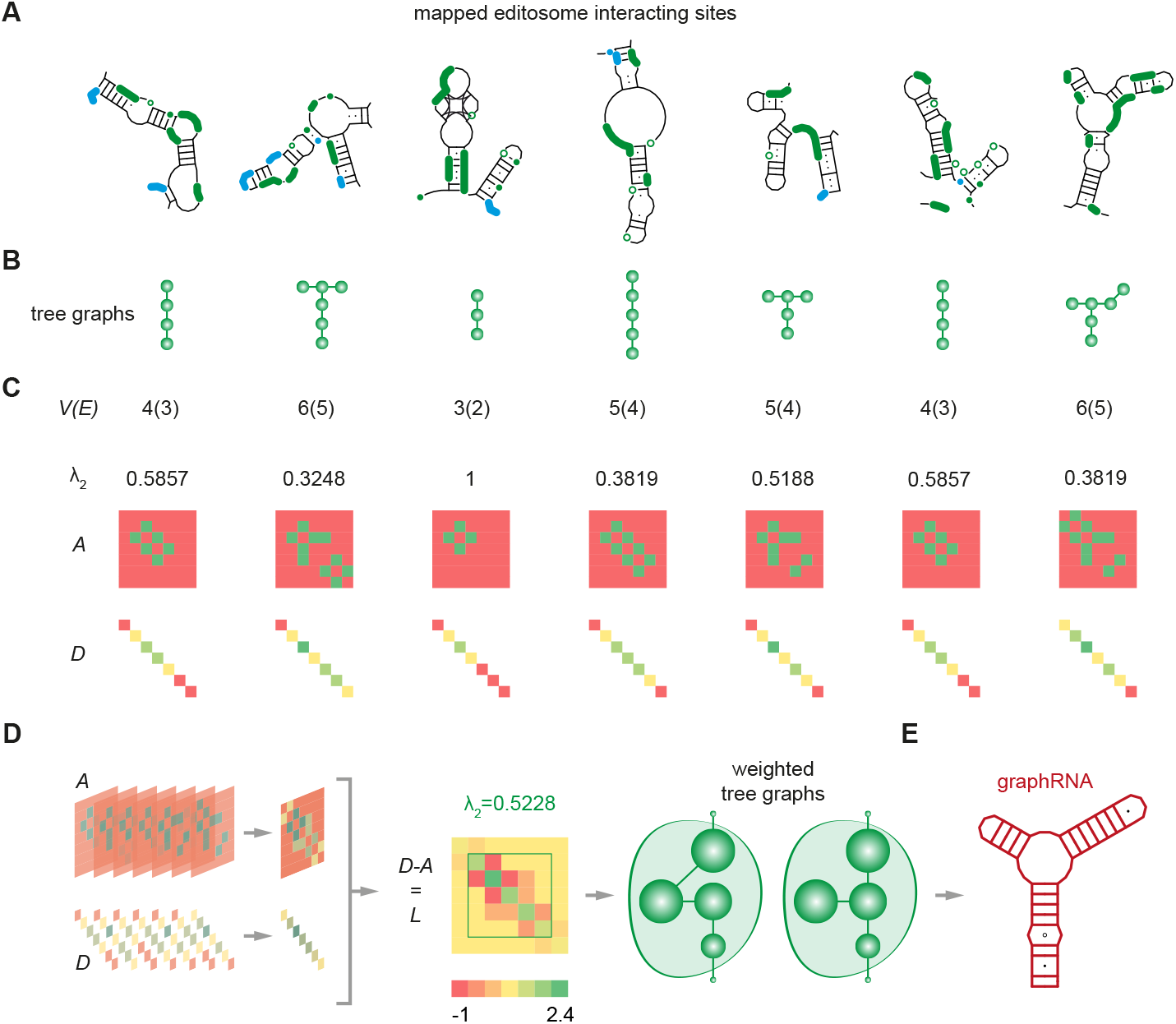
Graph theory-based RNA-motif analysis. (A) 2D-structures of the seven identified editosome interacting RNA-motifs in the RPS12-, ND3-, A6-, CO3- and ND7-pre-mRNAs. (B) Tree graph representations of the different RNA-motifs. Vertices (*V*)=green dots. Edges (*E*)=connecting lines. (C) Matrix representations of the different RNA-motifs: *A*=adjacency matrix, specifying the number of edges that connect pairs of graph vertices. *D*=square diagonal matrix detailing the connectivity of each vertex. λ_2_=2^nd^ Laplacian eigenvalue, enumerating the overall pattern of connectivity. (D) 3D-matrix stacks generated by concatenation of the individual *A*- and *D*-matrices. The stacks can be averaged along the 3^rd^-dimension to calculate mean *A*- and *D*-matrices. *L*=Laplacian matrix (*L*) defined as *L*=*D*-*A*. The resulting “averaged tree graph” is 4*V*(3*E*) with a λ_2_ eigenvalue of 0.5228. A statistical weighting of the individual vertices is indicated by the varying circle diameters. (E) Secondary structure outline of a representative 4*V*(3*E*)-RNA termed “graphRNA”. The molecule has a three-helix-junction topology and contains 2 GU-bp (filled dots) and a U/U-mismatch (open circle) to account for the complex folding architectures of the different editosome-interacting RNA-motifs.

### Design of a synthetic 4*V*(3*E*)-RNA

Based on the analysis above we designed a synthetic, 60nt long 4*V*(3*E*)-RNA, which we termed graphRNA (Fig. 6A). The molecule represents a mimic of the 7 identified pre-mRNA binding motifs and is characterized by helical segments 4bp, 8bp and 11bp in length. The three stems are connected through a multi-loop sequence and in order to generate structural plasticity we designed the shortest helix as an A:U-rich stem. Furthermore, we added a U:U mismatch and two G:U bp into the two other stems. GraphRNA is purine-rich with an R/Y-ratio of 1.3. It has a calculated ΔG of −0.36kcal/mol/nt and the majority of base pairs have a bp-probability ≥90% with a configurational entropy of 0.13. GraphRNA was chemically synthesized and after purification it was characterized by denaturing gel electrophoresis and temperature-dependent UV-spectroscopy (Fig. 6A). The RNA displays two melting transitions at 59°C and 73°C, which is consistent with a helical arrangement in which the 4bp helix stacks on top of the 8bp stem to form a quasi-continuous helix (Fig. 6B). A 3D-model of graphRNA is shown in Fig. 6B.

**Fig. 6.**
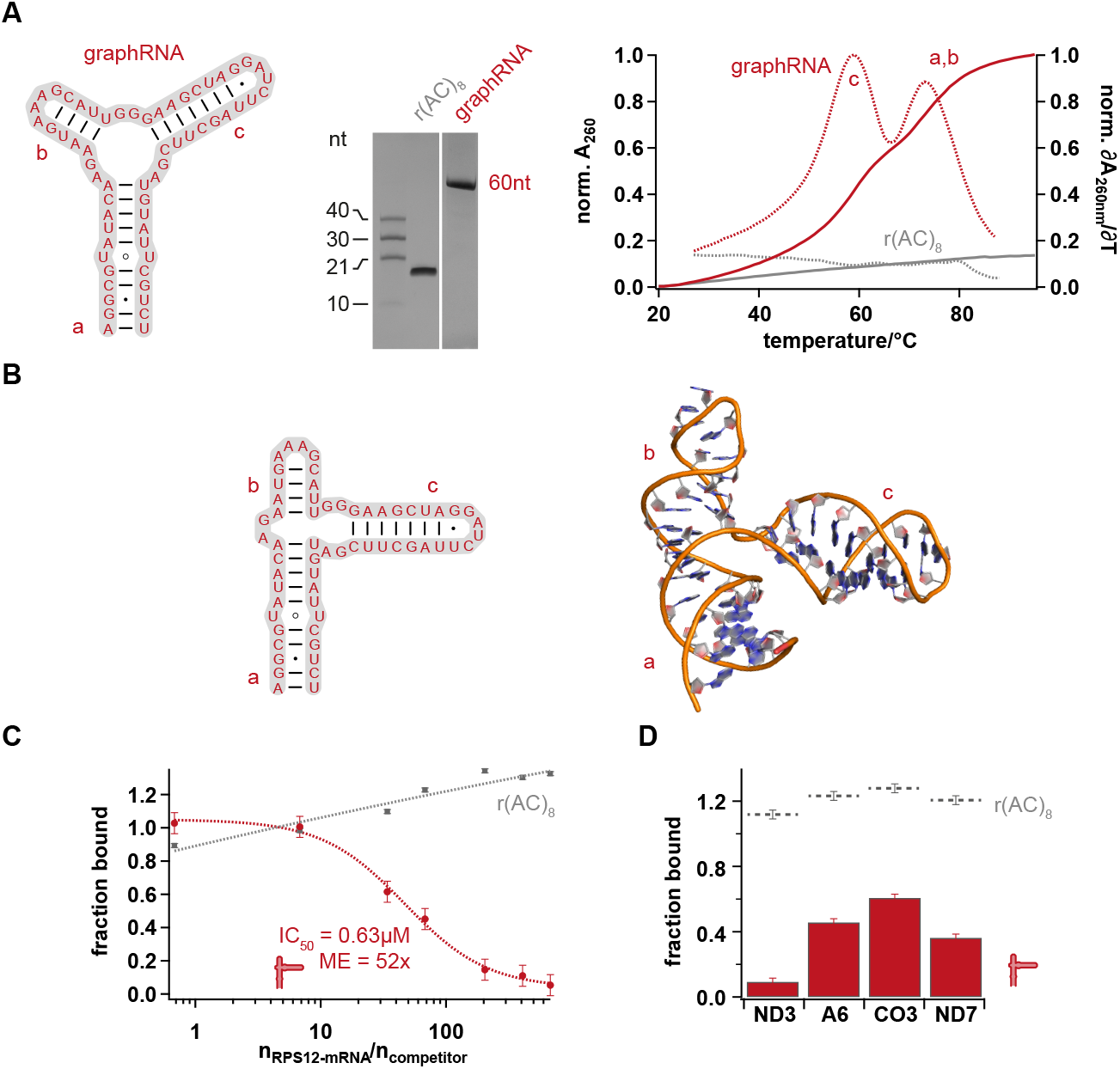
GraphRNA synthesis and binding competition for the editosomal RNA-binding site. (A) Left: 2D-structure and sequence of graphRNA with helical elements labeled a, b and c. Center: Gel electrophoretic analysis of chemically synthesized graphRNA in a urea-containing (8M) 10% (w/v) polyacrylamide gel in comparison to the single-stranded ribohexadecamer r(AC)_8_. The RNA-preparation shows the expected molecular length of 60nt and is ≥97% pure. Right: UV-melting (A_260_=f(T)) and 1^st^-derivative curves (δA_260_/δT)=f(T)) of graphRNA (red) and r(AC)_8_ (grey). GraphRNA exhibits two melting transitions at 59°C and 73°C in line with a stacked helical arrangement of helix b on top of helix a (B) as predicted by the 3D-model in (B). (C) Binding competition of (^32^P)-radiolabeled pre-mRNA RPS12 to the editosome by increasing concentrations of graphRNA. Half-maximal displacement occurs at a 52-fold molar excess (ME) of graphRNA over RPS12-transcript (arrow). The unstructured r(AC)_8_-oligoribonucleotide is not able to compete for the editosome binding site even at a ≥680-fold molar excess. (D) At a 500-fold molar excess of graphRNA, 90% of the ND3-transcript, 60% of the ND7 pre-mRNA, 45% of the A6-mRNA and 40% of the CO3-transcript are displaced from the editosome despite the fact that the synthetic RNA-molecule is 6-16-times smaller than the natural transcripts. Error bars are relative errors in percent.

### Binding of graphRNA to the editosome

The ability of graphRNA to bind to the editosome was tested in binding competition experiments. For that we synthesized (^32^P)-labeled preparations of all 5 pre-mRNAs (RPS12, ND3, ND7, A6, CO3). The radiolabeled transcripts were mixed with increasing concentrations of graphRNA and incubated with editosomes as the binding target. Fig. 6C shows a representative binding competition experiment using the RPS12 pre-mRNA as an example. While the single-stranded, hexadecameric control RNA r(AC)_8_ is unable to compete with the RPS12-transcript for the editosome binding site, graphRNA acts as a competitor. The competition curve has a sigmoidal shape with a linear range spanning one order of magnitude. Half-maximal competition is achieved at a 52-fold molar excess of graphRNA over RPS12-RNA. Similarly graphRNA competes with all other *T. brucei* pre-mRNA species (ND3, A6, CO3, ND7). Inhibition efficiencies range from 90% for the ND3-transcript to 40% for the pre-edited CO3 mRNA (Fig. 6D).

### GraphRNA inhibits *in vitro* RNA-editing

The result above tempted us to explore whether graphRNA not only interferes with the ability of the editosome to bind pre-mRNAs but also with the catalytic activity of the complex. For that we made use of the recently established U-deletion *in vitro* RNA editing assay (Del Campo et al., 2020; Leeder et al., 2021). The assay utilizes a synthetic, fluorescently labelled pre-mRNA/gRNA-hybrid RNA, which is edited by the site-specific deletion of 4 U-nt (Fig. 7A). The RNA-input molecule, the product of the catalytic conversion and all intermediates and side-products of the reaction can be traced in the assay and are separated with nucleotide resolution by capillary electrophoresis. As shown in Fig. 7B, graphRNA inhibits the U-deletion *in vitro* reaction in a concentration-dependent fashion. The catalytic conversion is half-maximally stalled at a concentration of 0.23μM, which corresponds to a 70-fold molar excess of graphRNA over the input pre-mRNA/gRNA hybrid (Fig. 7D). As before, the single stranded r(AC)_8_ control RNA is not able to inhibit the U-deletion reaction even at a 2500-fold molar excess (Fig. 7C,D). This suggests that graphRNA binds to the editosome near or at the catalytic reaction center of the complex.

**Fig. 7.**
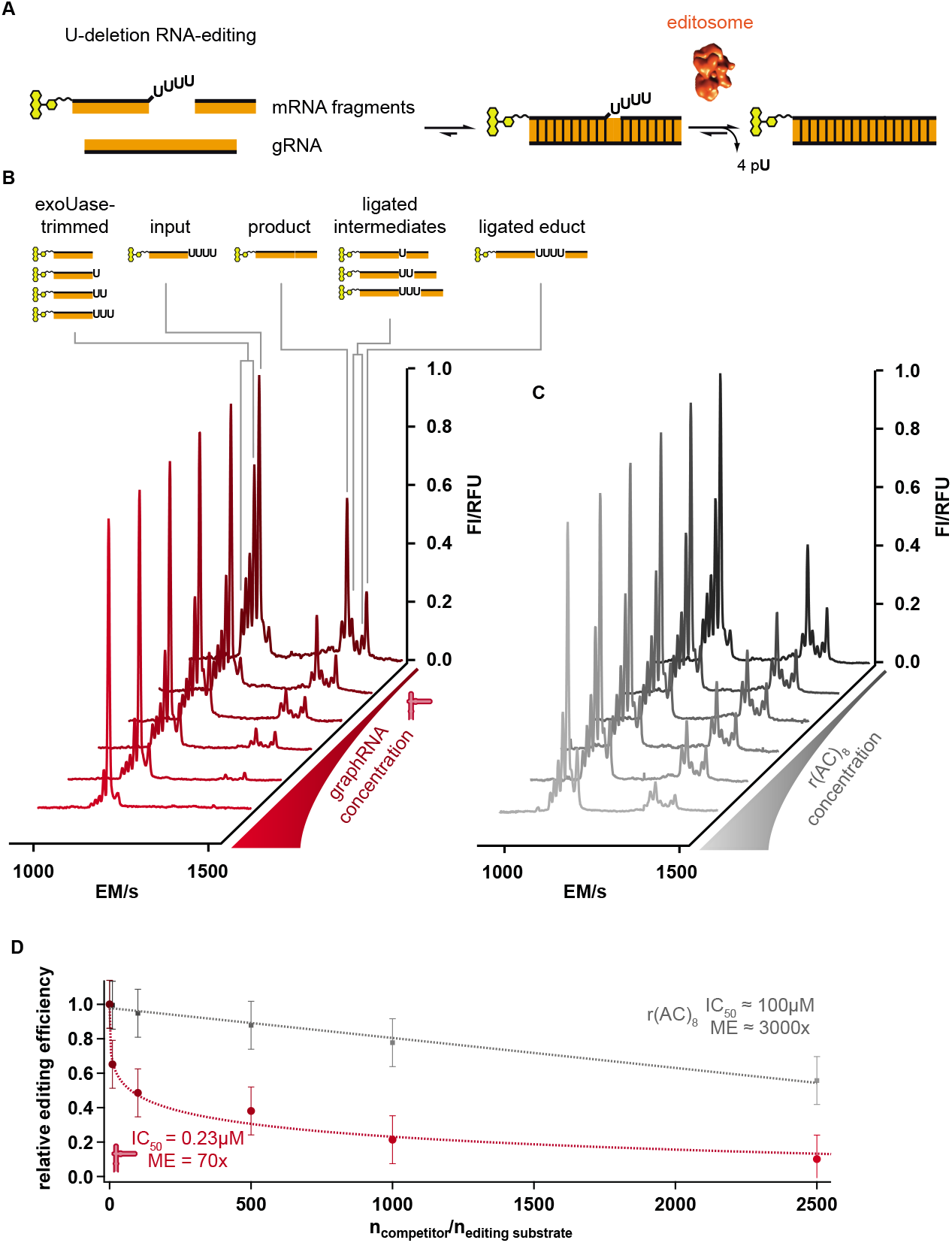
Inhibition of RNA-editing by graphRNA. (A) Diagram of the fluorescence-based U-deletion *in vitro* RNA-editing assay (Del Campo et al., 2020). The test system uses short oligoribonucleotides that mimic 2 pre-mRNA fragments and a complementary gRNA-molecule. Phosphate-ribose backbones are in black and nucleobases in orange. The 5’ pre-mRNA fragment is fluorophore-derivatized (yellow) to enable detection by laser-induced fluorescence. U-nucleotides to be deleted are depicted as capital letters. Formation of the trimolecular pre-mRNA/gRNA-substrate is driven by Watson-Crick base pairing (vertical bars) and the addition of editosomes (red sphere) initiates the deletion of 4 U-nt (pU). (B) Capillary electrophoresis profiles (red traces) of a representative U-deletion inhibition experiment using increasing concentrations of graphRNA (back to front). The assay resolves next to the +4U input-RNA, the edited −4U product as well as all intermediates and by-products. Cartoons of the different RNA-species are depicted on top of the individual peaks. FI= fluorescence intensity. RFU=relative fluorescense unit. EM=electrophoretic migration time. (C) CE-profiles (grey traces) of a U-deletion inhibition experiment using increasing concentrations of the single stranded control oligoribonucleotide r(AC)_8_ (back to front). (D) Quantitative representation of the data shown in (B) and (C). Plotted is the relative RNA-editing efficiency in relation to the molar ratio of competitor RNA to editing substrate RNA (ngraphRNA/nediting substrate). Error bars are relative errors in percent. ME=molar excess.

### Mono- and bimolecular graphRNAs are recognized by the editosome

Intriguingly, the three-way junction topology of graphRNA has a natural equivalent in the form of the gRNA/pre-mRNA hybrid RNAs of the editing reaction (Fig. 8A). Guide RNAs provide the sequence information for the U-insertion/deletion reaction by hybridizing to cognate pre-mRNAs and the resulting RNA-hybrids adopt archetypical 3-helix junction geometries (Lescoute and Westhof, 2006; De la Peña et al., 2009). Guide RNAs are with only one exception trans-acting RNAs (Golden and Hajduk, 2005). As a consequence, gRNA/pre-mRNA-hybrid RNAs are bimolecular assemblies, which tempted us to synthesize a bimolecular version of graphRNA. The molecule was termed graphRNA-split (Fig. 8A). GraphRNA-split was formed by hybridization of 2 oligoribonucleotides (graphRNA_split-5’ and graphRNA_split-3’) 34nt and 26nt in length. Annealing of the hybrid was confirmed by electrophoresis in non-denaturing polyacrylamide gels (Fig. 8B) and by temperature-dependent UV-hyperchromicity measurements (Fig. 8C). Identical to its monomolecular cousin, graphRNA-split it is able to inhibit *in vitro* U-deletion editing in a concentration-dependent fashion (Fig. 8D). Product formation is half-maximal suppressed at a concentration of 20nM, which represents a 6-fold molar excess of graphRNA-split over the input pre-mRNA/gRNA hybrid RNA (Fig. 8E). Thus, graphRNA-split is about 10-fold more efficient in inhibiting RNA-editing than the monomolecular version of the synthetic 4*V*(3*E*)-RNA.

**Fig. 8.**
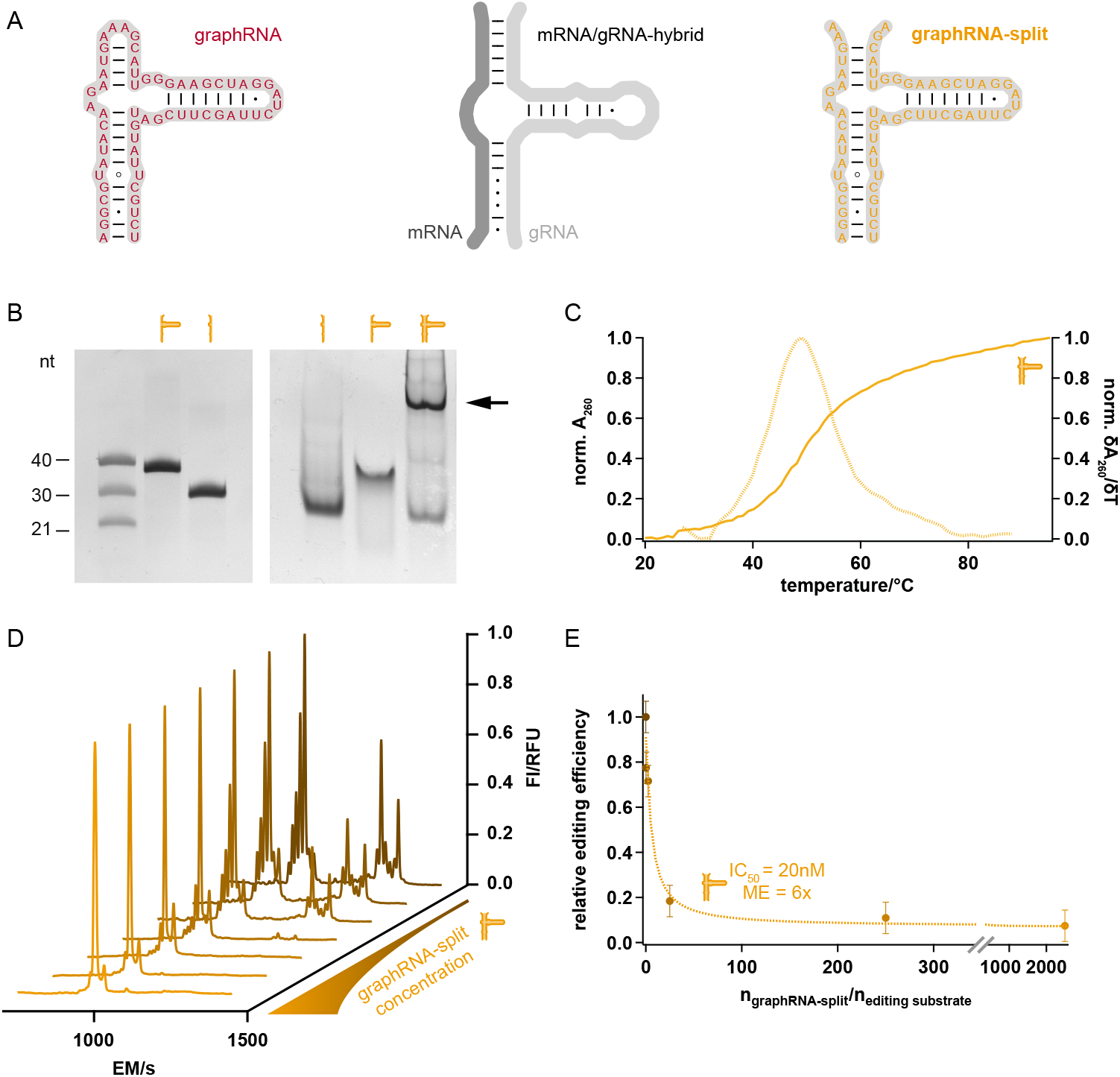
Inhibition of RNA-editing by graphRNA-split. (A) Secondary structure of graphRNA next to the 2D-structure of an archetypical gRNA/mRNA-hybrid RNA and graphRNA-split. (B) Left: Gel electrophoretic analysis of the chemically synthesized 5’- and 3’-fragments of graphRNA-split analyzed in a urea-containing 18% (w/v) polyacrylamide gel. Right: Annealing of the 5’- and 3’-oligoribonucleotides to form graphRNA-split analyzed in a non-denaturing 18% (w/v) polyacrylamide gel. The bimolecular reaction product is marked by an arrow. (C) UV-melting (A_260_=f(T)) and 1^st^-derivative curves (δA_260_/δT)=f(T)) of graphRNA-split. The molecule shows a melting transitions at 49°C. (D) Capillary electrophoresis profiles of a representative U-deletion inhibition experiment using increasing concentrations of graphRNA-split (brown to yellow). Cartoons of the different RNA-species (peaks) are shown in Fig. 7B. FI= fluorescence intensity. RFU=relative fluorescense unit. EM=electrophoretic migration time. (E) Quantitative representation of the data shown in (D). Plotted is the relative RNA-editing efficiency in relation to the molar ratio of graphRNA-split to editing substrate RNA (n_graphRNA-split_/n_editing substrate_). U-deletion RNA-editing is inhibited with an IC_50_ of 20nm equivalent to a 6-fold molar excess (ME). Error bars are relative errors in percent.

## Discussion

The U-nucleotide specific RNA-editing reaction in the mitochondria of kinetoplastid organisms such as African trypanosomes has been identified as an inherently noisy process. This is based on the observation that steady state isolates of mitochondrial RNA contain next to pre-edited mRNAs large quantities of partially edited and mis-edited transcripts (reviewed in Zimmer et al., 2018). As a consequence, editosomes, the protein complexes that catalyze the editing reaction, must be able to recognize a pool of RNA-molecules that not only is highly divergent in primary sequence and nucleotide length but also in its U-content. Moreover, both the primary sequences and the U-content constantly change as the reaction advances, which makes the binding of substrate RNAs into the single RNA-binding site of the editosome (Böhm et al., 2012) a moving-target-problem. How editosomes untangle the structural multiplicity and dynamic disorder in the RNA-substrate landscape is unknown.

In order to unravel the RNA-recognition characteristics of the editosome we determined the RNA-interaction sites of five *T. brucei* mitochondrial pre-mRNAs experimentally with nucleotide resolution. We used the differential RNA-solvent accessibility between the free and editosome-bound folding states as a read-out and probed the solvent accessibility by high throughput hydroxyl radical footprinting as established by Weeks and colleagues (Duncan and Weeks, 2010a,b). Editosomes have been shown to bind pre-edited mRNAs with nanomolar affinity (Böhm et al., 2012) and thus, as anticipated, binding of the 800kDa protein complex generated discernable footprints in all pre-mRNAs. The majority of solvent-protected nucleotides cluster in and around defined secondary structure elements in all transcripts, which vary in length between 40 and 60nt. In addition, we identified RNA-sequence stretches that become more solvent exposed upon editosome binding. This demonstrates that binding of the protein complex induces a partial refolding in the different pre-mRNAs in line with the documented RNA-chaperone activity of the editosome (Böhm et al., 2012; Leeder et al., 2016b; Voigt et al., 2018). Thus, as for many RNA/protein complexes the formation of pre-mRNA/editosome complexes involves an RNA-motif recognition-step and an RNA-structure refolding-step (Balcerak et al., 2019; Jankowsky and Harris, 2015). However, unlike in other RNA/protein complexes the identified editosome-interacting RNA-motifs are similar but not identical. All attempts to identify a shared nucleotide sequence or common 2D-fold in the solvent-protected RNA-sequences failed. This suggests that the shared characteristics of the identified RNA-motifs are somehow concealed on the nucleotide level and as a consequence require a different analytical and conceptual framework. Since RNA secondary structures are essentially 2D-networks, they can be represented as 2-dimensional, coarse-grained graphs (Schlick, 2018; Gan et al., 2003). By using only vertices and edges as descriptors we transformed all solvent-protected RNA-motifs into planar, tree-like graph objects, which reduced the individual RNA-motifs to their connectivity attributes. This in turn enabled a mathematical comparison of the graphs, ultimately permitting the description of a shared tree-graph consisting of 4 vertices and 3 edges (4*V*(3*E*)). Every identified solvent-protected RNA-motif is a representative of this shared graph object, demonstrating that the RNA-recognition step of the editosome is characterized by a high degree of plasticity or “fuzziness” despite the fact that it is of high affinity (Shen et al., 2018; Järvelin et al., 2016). Obviously the editosome does not recognize a precisely folded mRNA-motif, instead it allows for conformational ambiguity similar to what has been described for “fuzzy” protein/protein complexes (Sharma et al., 2015).

As a proof-of-concept we re-transformed the identified consensus graph back into an RNA-molecule that matches all identified graph criteria and as a consequence should function as a synthetic, editosome-binding RNA-motif. Of the different RNA-folds that are consistent with the underlying graph we synthesized an RNA of mediocre thermodynamic stability with a length of 60nt. The molecule was termed graphRNA. As predicted, graphRNA binds to the editosome and competes with all five pre-mRNAs for binding to the complex despite the fact that it is significantly smaller. Furthermore, our data suggest that graphRNA binds near or at the catalytic center of the editosome since the synthetic RNA also inhibits U-deletion RNA-editing *in vitro*. Interestingly, graphRNA adopts a prototypical 3-helix-junction fold and in that it resembles the Y-shaped structure of a gRNA/pre-mRNA hybrid. Guide RNAs are central to the editing process by functioning as templates in the reaction. They execute their role by base pairing to the pre- and partially edited mRNAs proximal to an editing site and since in the majority of cases gRNA/mRNA-hybrids are bimolecular assemblies, we also analyzed a bimolecular or “split” version of graphRNA. GraphRNA-split behaves identical to its monomolecular cousin and inhibits *in vitro* U-deletion editing in a concentration-dependent fashion. This demonstrates that an unrelated, synthetic RNA-molecule can bind to the editosome as long as it fulfills the fuzzy, 4*V*(3*E*)-RNA folding criteria. Importantly, the identified folding constraints include the general topology of gRNA/mRNA-hybrid RNAs and as demonstrated before, editosomes are capable of binding both, mono- and bimolecular gRNA/mRNA hybrids. Based on these data it is tempting to speculate that the editosome has evolved to preferentially recognize the large folding space of the gRNA/mRNA hybrid RNAs. At steady state, literally hundreds of hybrid RNAs are present in the mitochondria of African trypanosomes, which adopt similar but non-identical 3-way-junction folds. Thus, only a non-binary, flexible or fuzzy RNA-recognition principle is able to respond to the highly diverse landscape of editing substrates (Fig. 9A,B), which is further increased by the large pool of partially edited and misedited mRNAs (Zimmer et al., 2018).

**Fig. 9.**
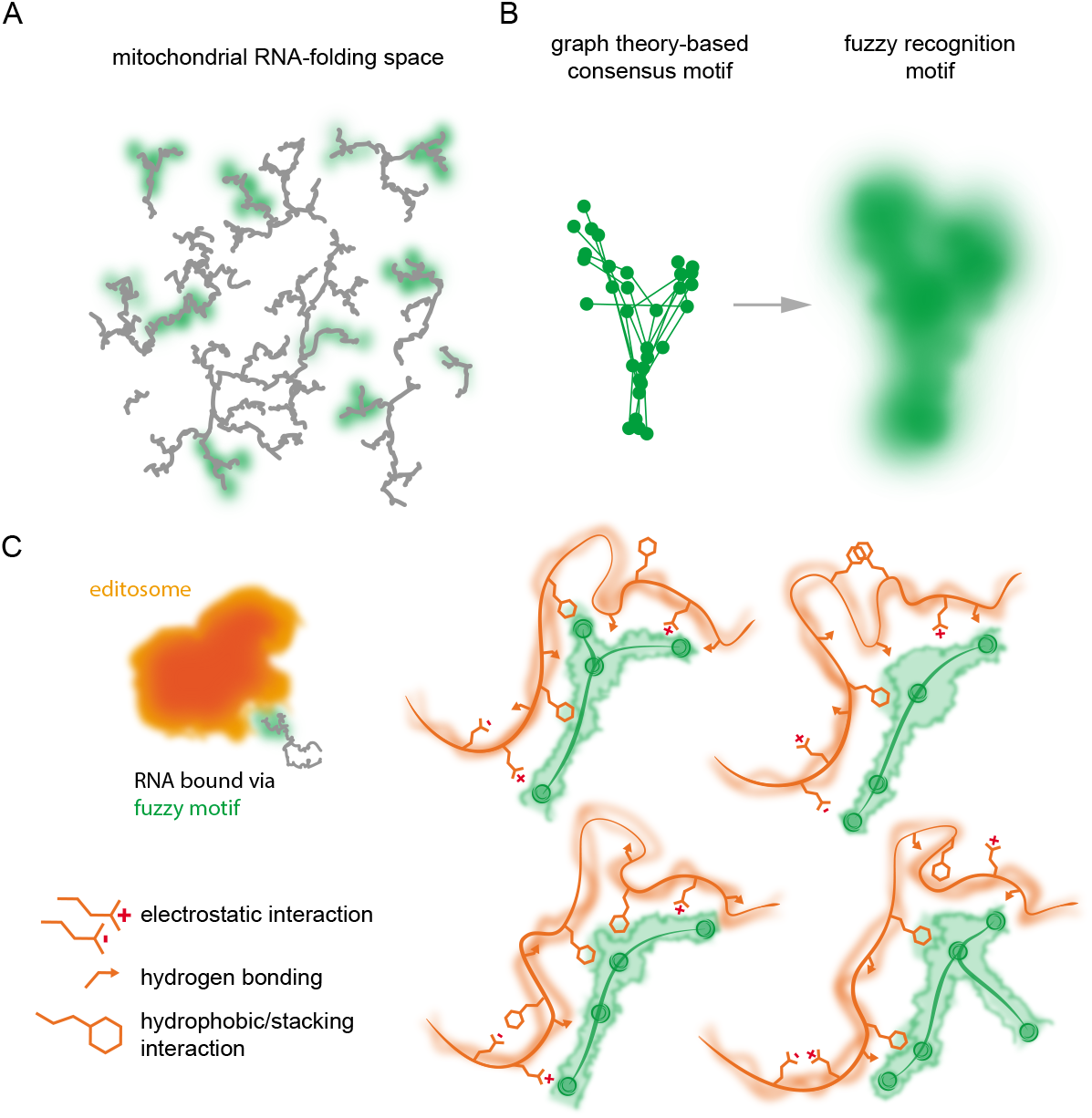
Fuzzy RNA-recognition by the *T. brucei* editosome. (A) Schematic representation of the convoluted RNA-folding landscape of the *T. brucei* mitochondrial transcriptome. RNA-species that are substrates of the RNA-editing reaction are characterized by one or more editosome-binding motif (green blurs), which all fit the graph theory-derived 4*V*(3*E*)-criterion (B). The multitude of similar but non-identical RNA-motifs can be viewed as a fuzzy-type recognition mode, which enables the editosome to interact with a highly divergent/disordered RNA-folding space. (C) We hypothesize that the RNA/protein interface relies on multiple physicochemical interaction principles including strong and week forces with a high degree of conformational entropy. This allows for structural flexibility and includes the capacity to self-modulate and self-adjust.

Which of the editosomal protein components is able to execute a fuzzy RNA-binding functionality? While only 4 of the roughly 20 editosomal proteins contain canonical RNA-binding domains (Czerwoniec et al., 2015) recent RNA-interactome screening experiments in human cells revealed that many RNA-binding proteins (55%) lack archetypical RNA-binding domains. Instead, they frequently harbor intrinsically disordered protein (IDP)-domains (Balcerak et al., 2019). IDP’s bind RNA with high specificity by providing large complementary binding surfaces, which depend on structural flexibility and a coupling of the folding and binding steps. Importantly, editosomal proteins are exceptionally rich in IDP-domains. Nearly 30% of the amino acid sequences are disordered, which is higher than the average of the human proteome (22%) (Czerwoniec et al., 2015). Particularly the six oligonucleotide/oligosaccharide-binding (OB)-fold proteins and here specifically the editosomal core proteins KREPA1, KREPA2 and KREPA3 are significantly more disordered than the average of all other editosomal proteins. As demonstrated by Voigt et al., 2018 five of the six OB-fold proteins execute RNA-chaperone activity to partially resolve the highly folded pre-edited substrate mRNAs (Leeder et al., 2016a,b). The activity is connected to the IDP-domains of the proteins, which have been proposed to be localized on the surface of the high-molecular mass complex (Voigt et al., 2018). Thus, we hypothesize that the demonstrated fuzzy RNA-binding property of the *T. brucei* editosome is executed by the IDP-domains of some or all of the editosomal OB-fold proteins (Supplementary Fig 5). This is supported by the fact that ID-proteins have been shown to sample a broad range of dynamic structural changes upon interacting with partner molecules. This includes the formation of fuzzy protein complexes, which are characterized by retaining a disordered folding state during complex formation (reviewed in Sharma et al., 2015). Prominent examples are the assembly of RNP-stress granules (Protter et al., 2018), protein liquid-liquid phase separation (Wu and Fuxreiter, 2016; Boeynaems et al., 2018) and RNA/capsid-protein interactions in viral systems (Ivanyi-Nagy et al., 2008; Byk and Gamarnik, 2016). We propose that the identified fuzzy RNA-motifs in the *T. brucei* pre-mRNAs mirror the spatially disordered folding characteristics of the editosomal IDP-domains, which enable the protein complex to interact with multiple mitochondrial RNA-species and multiple RNA-recognition motifs simultaneously. As a result, the editosome is able to address the highly divergent mitochondrial RNA-folding space including all pre-, partially and mis-edited mRNAs in addition to the large pool of gRNAs. Furthermore, we predict that the RNA-recognition step of the editosome follows a disorder-to-disorder transition in which both, the protein- and the RNA-contact sites remain in a state of high conformational entropy (Miskei et al., 2020) (Fig. 9C). Such a setup is characterized by a high degree of structural flexibility and includes the capacity to self-modulate and to self-adjust. By relying on multiple physicochemical interaction principles involving attractive and repulsive forces (Gruet et al., 2016) a fuzzy molecular interaction is not only able to dynamically modulate the binding reaction, it is also able to respond to environmental fluctuations (Miskei et al., 2020; Horvath et al., 2020; Fuxreiter, 2020a,b). This phenomenon might be responsible for the documented ability of the *T. brucei* editosome to edit mitochondrial mRNAs in both parasite life cycle stages at dissimilar redox and temperature conditions (27°C in insect-stage trypanosomes *versus* 37°C in bloodstream-stage parasites).

Lastly, we predict that many more non-binary *i.e*. fuzzy RNA/protein interactions exist. Especially in situations in which proteins or protein complexes are required to respond to changing cellular conditions by scanning large ensembles of different RNA-molecules (Järvelin et al., 2016). As such, our finding represents a logical extension of the fuzzy protein theory for disordered proteins (Fuxreiter, 2020) by merging the fuzziness and frustration concepts in the energy landscape of proteins with that of RNA-molecules (Gianni et al., 2021). As much as frustration in fuzzy protein/protein complexes causes a multiplicity of specific interactions (Freiberger et al., 2021; Ferreiro et al., 2014) it is conceivable that frustration in RNA/protein complexes is responsible for the observed non-binary pre-mRNA-binding characteristics of the *T. brucei* editosome.

## Methods

### Oligonucleotide synthesis

DNA- and RNA-oligonucleotides were synthesized by automated solidphase synthesis using controlled pore glass (CPG)-beads and 2-cyanoethyl- (DNA) or 2’-O-(tert-butyl)dimethylsilyl (TBDMS)-protected (RNA) phosphoramidites. Fluorophore modifications (6-carboxy-fluorescein=FAM, 5’-dichloro-dimethoxy-fluorescein=JOE and 5-Carboxytetramethyl-rhodamine= TAMRA) were introduced post-synthetically using a 5’-terminal C6-aminolinker, which were conjugated by EDC (1-Ethyl-3-(3-dimethylaminopropyl)carbodiimide)-mediated coupling. Similarly, 3’-end NH2-modifications were introduced via a hexamethylene-aminolinker. Base-protecting groups were removed using NH4OH/EtOH (3:1) at RT and 2’-silyl protecting groups were removed using neat triethylamine trihydrofluoride. All oligonucleotides were HPLC-purified, analyzed by mass-spectrometry and further scrutinized in denaturing polyacrylamide gels. Oligonucleotide concentrations were derived from UV-absorbency measurements at 260nm (A_260_) using the molar extinction coefficients (ε in L/mol x cm) listed below. The following DNA- and RNA-sequences were synthesized:

Reverse transcriptase (RT)-primer:

T3_rev FAM- or JOE-(CH_2_)_6_-AATTAACCCTCACTAAAGGGAAC (292300)
CO3_rev_317-344 JOE-(CH_2_)_6_-GTATTCCTTTGCCCAAAAACCCCTTTG (300900)
CO3_rev_225-196 FAM- or JOE-(CH_2_)_6_-CCTCAAAACCTCCTCTCAAAACAAACTCTT (345500)
CO3_rev_381-358 FAM- or JOE-(CH_2_)_6_-AAAACCCCTCCAAAAACCCTTCTC (278500)
ND7_rev_356-334 FAM- or JOE-(CH_2_)_6_-TTTCAAAATCTTTCGCTTGCCG (247900)
ND7_rev_452-428 FAM- or JOE-(CH_2_)_6_-AAAGCCTGCTCGCCCCAAAACTTCT (285300)
ND7_rev_601-574 FAM- or JOE-(CH_2_)_6_-CCTTCACAAAAATCAAAAATCCTTCGAC (340900)
ND7_rev_868-840 FAM- or JOE-(CH_2_)_6_-AAACAATCCTCAATAACCAAAAAATAACC (379500)
graphRNA AGGCGUAUACAAGAAUGAAAGCAUUGGGAAGCUAGGAUCUUAGCUUCGAUGUAUUCGUCU (700900)
graphRNA_split-5′ AGGCGUAUACAAGAAUGAAAGCAUUGGGAAGCUA (422600)
graphRNA_split-3′ GGAUCUUAGCUUCGAUGUAUUCGUCU (278300)
r(AC)_8__ssRNA ACACACACACACACAC (182400)
U-deletion editing substrate RNAs:

5′Cl22 FAM-(CH_2_)_6_-GGAAAGGGAAAGUUGUGAUUUU (292460)
3′Cl15 pGCGAGUUAUAGAAUA-(CH_2_)_6_-NH2 (186600)
gRNA_del GGUUCUAUAACUCGCUCACAACUUUCCCUUUCC (335400)
U-insertion editing substrate RNAs:

5′Cl18 TAMRA-(CH_2_)_6_-GGAAGUAUGAGACGUAGG (247200)
3′Cl13 pAUUGGAGUUAUAG-(CH_2_)_6_-NH2 (154400)
gRNA_ins CUAUAACUCCGAUAAACCUACGUCUCAAUACUUCC (359400)

### Pre-edited mRNA synthesis

Pre-edited mRNAs (A6, ND7, RPS12, COIII, ND3) were synthesized by run-off *in vitro* transcription from linearized plasmid DNA-templates. Typically, reactions were performed in 0.1mL in 40mM Tris-HCl pH7.9, 6mM MgCl_2_, 2mM spermidine, 10mM DTT containing 3μg linearized DNA-template, 1.5mM of each ATP, CTP, GTP, UTP, 40U Ribolock RNase inhibitor (Thermofischer Scientific) and 100U T7-RNA polymerase. Samples were incubated for 2h at 37°C and stopped by DNaseI digestion (15min, 37°C) followed by phenol extraction. Non-incorporated ribonucleotides were removed by size exclusion chromatography. RNA-concentrations were calculated from UV-absorbency measurements at 260nm (A_260_). The integrity of the different pre-mRNA preparations was analyzed in 6% (w/v) urea-containing (8M) polyacrylamide gels. Radiolabeled (^32^P) pre-mRNA preparations were synthesized in 0.02mL reactions as described above using 1μg linearized DNA-template, 6μM α-(^32^P)-UTP (400Ci/mmol), 12μM UTP and 50U T7-RNA polymerase.

### Editosome purification

Editosomes were purified from insect-stage African trypanosomes (*Trypanosoma brucei*) grown at 27°C in SDM-79 medium in the presence of 10% (v/v) fetal calf serum (Brun and Schönenberger, 1979). The complexes were isolated by tandem-affinity purification (TAP) (Gerace and Moazed, 2015) using transgenic *T. brucei* cell lines that conditionally express TAP-tagged versions of the editosomal proteins KREPA3, KREPA4 or KRET2 (Golas et al. 2009, Kala et al., 2012; Ringpis et al., 2010). Routinely about 6×10^11^ parasite cells were lysed and further processed using IgG- and calmodulin-affinity chromatography resins. The protein composition of the isolates was analyzed in SDS-containing polyacrylamide gels and the identity of the different proteins was determined by mass spectrometry. The integrity of the complexes was scrutinized by Atomic Force Microscopy (AFM) (Böhm et al., 2012) and the RNA-editing activity of the isolates was tested for both, the U-insertion and the U-deletion reaction (Del Campo et al., 2020; Leeder et al., 2021).

### Hydroxyl radical footprinting (HRP)

*In vitro* transcribed pre-edited mRNAs (3pmol) were refolded as described in Leeder et al., 2016a. Editosomes in editing buffer (EB = 20mM HEPES pH7.5, 30mM KCl, 10mM MgCl_2_, 0.5mM DTT, 0.5mM ATP) were added at a molar RNA: editosome ratio of 1:2 and were allowed to bind for 20min at 27°C. Note: the reaction time is ?5-times the half-life (t1/2) of pre-mRNA/editosome complexes (Katari et al., 2013) and 27°C represents the optimal temperature for the U-insertion/U-deletion reaction. Control samples contained EB only. Hydroxyl radicals were generated by the sequential addition of 265μM (NH_4_)Fe(SO_4_)_2_, 176μM EDTA, 0.007 % (v/v) H_2_O_2_ and 4.25mM Na-L-ascorbate (C_6_H_7_O_6_Na) in a total volume of 35μL up to 70μL (Fenton, 1894; Duncan and Weeks, 2010a,b). Samples were incubated for 45sec at 27°C, which corresponds to single-hit conditions within a 300-400nt sequence window. Reactions were terminated by EtOH-precipitation. RNA-precipitates were washed (70% (v/v) EtOH), resuspended in 10mM Tris-HCl pH7.5, 1mM EDTA (TE)-buffer followed by phenol-CHCl3-isoamylalcohol (25:24:1) extraction and desalting by size exclusion chromatography (Bio-Gel^®^ P6-resin). OH-radical induced phosphodiester cleavage sites were identified by abortive reverse transcription using fluorophore-substituted DNA-primer molecules as described in Leeder et al., 2016a. cDNA-fragments were resolved by automated capillary electrophoresis (Leeder et al., 2016a; Leeder and Göringer 2020) and raw electropherograms were processed using ShapeFinder (Vasa et al., 2008). HRP-data were statistically analyzed using custom Python scripts with the help of forgi 2.0 (Thiel et al., 2019).

### Editosome-binding competition assay

Binding competition experiments were performed in 35μL EB containing 12nM (^32^P)-labeled pre-edited RPS12-, ND3-, A6-, CO3- or ND7-mRNA (50.000cpm) and increasing amounts (up to a 680-fold molar excess) of unlabeled graphRNA or the single-stranded control RNA r(AC)_8_. Reactions were started by the addition of editosomes (15nM) and were incubated for 30min at 27°C. Samples were filtered through nitrocellulose (NC)-filter membranes (Merck Millipore HAWP 0025) at a rate of 7.5mL/min. NC-filters were washed with 30 reaction volumes (1mL) EB, dried and quantified by liquid scintillation counting.

### Inhibition of RNA-editing

Trimolecular U-deletion pre-mRNA/gRNA hybrid RNAs were formed by combining equimolar amounts (0.1μM) of the pre-mRNA mimicking RNA-oligonucleotides 5’Cl22 and 3’Cl15 with guide RNA gRNA_del. Samples were denatured for 1min at 75°C followed by cooling to 27°C at a rate of 0.2°C/sec. Annealed pre-mRNA/gRNA-hybrids (100fmol) in 30μL 20mM HEPES-KOH pH7.5, 30mM KCl, 10mM MgCl_2_, 3mM DTT, 0.5mM ATP, 60μM UTP were mixed with a molar excess of graphRNA (10- to 5000-fold), graphRNA-split (0.25- to 5000-fold) or with r(AC)_8_ (10- to 5000-fold) as a single-stranded control oligoribonucleotide. Reactions were started by the addition of 2.5nM editosomes. After incubation for 30min at 27°C reactions were stopped by phenol extraction and RNAs were recovered by EtOH-precipitation and analyzed by CE. Raw electropherograms were baseline corrected followed by peak integration (Del Campo et al., 2020; Leeder et al., 2021). Relative RNA-editing efficiencies were calculated as the ratio of edited product-RNA over input-RNA and were normalized to a control reaction without competitor RNA. Data were plotted as a function of the competitor-RNA/input-RNA ratio and the resulting dose-response curves were fitted to the Hill-Langmuir equation (θ_ed_= 1/1+(K_a_/[RNA])^n^) to derive half-maximal inhibitory concentrations (IC_50_) (θ_ed_=fraction of RNA-bound editosomes, [RNA]=free RNA concentration, K_a_=RNA-concentration at half-maximal occupation, n=Hill-coefficient).

### RNA-As-Graphs (RAG) analysis and RNA-modeling

Solvent-protected RNA-motifs defined as 2Dstructure elements displaying clustered sequence stretches with negative ΔHRP-values (over a 3nt window) below the 25^th^ percentile were analyzed using the RNA-As-Graphs (RAG) resource developed by Schlick and colleagues (Fera et al., 2004; Gan et al., 2004). RAG converts RNA 2Dstructures into coarse-grained 2D-tree graphs and classifies them by their vertex (*V*) number and eigenvalue spectrum. The 2D-structure of graphRNA was designed using ViennaRNA package 2.0 (Lorenz et al., 2011). The data were used as constraints in a coarse-grained replica exchange Monte Carlo (REMC)-molecular dynamics (MD) simulation using SimRNA (Boniecki et al., 2016). *In toto* 10 replicas of simulated annealing were generated with 1.3×10^6^ iterations each. The resulting trajectories were submitted to a clustering of the 1% lowest energy conformations and grouped with a root-mean-square deviation (RMSD)-threshold of 5Å.

## Acknowledgements

We thank Andreas Völker for trypanosome cell growth, Robert Knieß and Cristian Del Campo for editosomes preparations and Reza Salavati, Inna Aphasizheva and Ruslan Aphasizhev for transgenic trypanosome strains. This work was supported by grants from the German Research Foundation (DFG-GO516/8-1) and the Dr. Illing-Foundation for Molecular Chemistry to H.U.G.

## Author contributions

H.U.G. and W.-M.L. designed the study. W.-M.L. performed the experiments. Both authors interpreted the data and wrote the manuscript.

## Additional information

Supplementary information. The online version contains supplementary material.

