## Supplementary information for "Fuzzy RNA-recognition by the *Trypanosoma brucei* editosome"

Wolf-Matthias Leeder, H. Ulrich Göringer
Molecular Genetics, Technical University Darmstadt, 64287 Darmstadt, Germany

**Supplementary Figure 1. Characteristics of the pleomorphic structural landscape of *T. brucei* mitochondrial pre-mRNAs.** (A) Summary of the RNA-editing induced changes in nucleotide (nt) length, purine/pyrimidine (R/Y)-ratio and Gibbs free energy per nucleotide (ΔG/nt) for the mitochondrial pre-mRNAs ND3, ND8, RPS12, CR4, ND9, CR3, A6, Cyb, CO3, MURF2 and ND7. (B) Selection of 5 minimal free energy (MFE)-structures of the *T. brucei* CR4-transcript during its transition from a pre-edited mRNA (left) to a fully edited transcript (right). Pre-edited CR4-mRNA contains 3 GQ-elements (leaf-like structures in red), which are progressively resolved to generate a fully edited, GQ-free mRNA. (C) Next to the different MFE-structures numerous alternative 2D-folds exist in the thermodynamic ensemble as illustrated by the base pairing probability-matrices below the 5 CR4 MFE-folds shown in (B). The 2D-structures are composed of highly probable base pairs (red dots) as well as less probable base pairs in orange, green and blue.

**Supplementary Figure 2. Artistic rendering of the solvent-accessibility/solvent-inaccessibility concept to identify editosome/RNA-contact sites**. (A) Molecular model of the *T. brucei* 20S editosome (Voigt et al., 2018) based on the cryoEM-derived shape of the protein complex (Golas et al., 2009). (B) RNA-binding to the editosome generates solvent-accessible (brush stroke in blue) and solvent-inaccessible (green) RNA-surfaces, which can be mapped by hydroxyl radical footprinting (HRP). (C) Solvent-accessible RNA-domains are HRP-sensitive and solvent-inaccessible RNA-segments are HRP-protected.

**Supplementary Figure 3. The pre-mRNA/editosome interface has a finite size.** (A) Schematic drawings of 3 editosome/RNA-complexes involving a small-, medium- and large-size pre-mRNA (from left to right) bound to the single RNA-binding site of the complex (Böhm et al., 2012). RNA-molecules are depicted as 2D-silhouettes and editosomes are in red. The RNA/editosome interface is marked by a green line. (B) Plot of the number of highly HRP-protected nucleotides (defined as the $25^{th}$-percentile of negative ΔHRP-values - (IQR x 0.7) within the top 10% of negative ΔHRP-values) *versus* the total number of HRP-probed nucleotides in the RPS12-, ND3-, A6-, CO3- and ND7-transcripts. The data fit a saturation function suggesting that a finite number of roughly 60nt are involved in the editosome/RNA interaction. (C) Boxplot of the ΔHRP-values of the different pre-mRNAs plotted as a function of the total number of interrogated nucleotides (RPS12=336nt, ND3=376nt, A6=398nt, CO3=705nt, ND7=1057nt). The data show no transcript length-dependence.

**Supplementary Figure 4. Comparison of nucleotide-solvent accessibility and local nucleotide dynamic.** (A) Scatter-plot analysis of the HRP-solvent accessibility data with published SHAPE-modification data (Leeder et al., 2016b; Leeder et al., 2020) for the *T. brucei* pre-mRNAs RPS12, ND3, A6, CO3 and ND7 (left to right). SHAPE maps nucleotide dynamic and as evidenced by Pearson correlation coefficients ($p$) between 0.1 and 0.4 the nucleotide-solvent accessibility is not correlated with the flexibility of nucleotides. (B) Plots of the ΔHRP-data for the RPS12, ND3, A6, CO3 and ND7 pre-mRNAs (left to right) *versus* SHAPE-reactivity. With $p$-values between -0.2 and 0.01 the editosome-induced reactivity changes for OH-radicals are not biased towards flexible or inflexible nucleotides. HU=HRP-unit. SU=SHAPE-unit.

**Supplementary Figure 5. Fuzziness and order/disorder propensities of *T. brucei* editosomal OB-fold proteins.** FuzPred-analysis (Horvath et al., 2020; Miskei et al., 2020) of the *T. brucei* oligosaccharide/oligonucleotide binding (OB)-fold proteins KREPA1, KREPA2, KREPA3, KREPA4, KREPA5 and KREPA6. FuzPred predicts the binding mode landscape of intrinsically disordered proteins based on their amino acid sequences. (A) The probability for a disorder-to-disorder binding mode ($P_{DD}$) for each amino acid (Ai) in the different proteins is calculated over a 5-9 amino acid window in grey. Black traces: The chance of any Ai-residue to act in a context dependent fashion is referenced as the Shannon entropy ($S_{Ai}$) calculated from the probability of all possible binding modes. High $SA_i$-values (≥2.25) characterise fuzzy Ai-positions with a high variety of binding modes (green). (B) Plot of the context dependency expressed as the Shannon entropy ($S_{Ai}$) as a function of the disorder-to-disorder binding mode probability ($P_{DD}$) for all proteins. (C) The OB-fold proteins KREPA1 to KREPA6 are shown as white circles arbitrarily placed inside the cryoEM-outline of the *T. brucei* editosome (Golas et al., 2009). Circle sizes are scaled to the amino acid length of the different proteins. Sequence stretches ≥5 amino acids that are predicted to exert fuzzy DD-interactions are depicted as green circles. Predicted non-fuzzy DD-interactions are in grey.

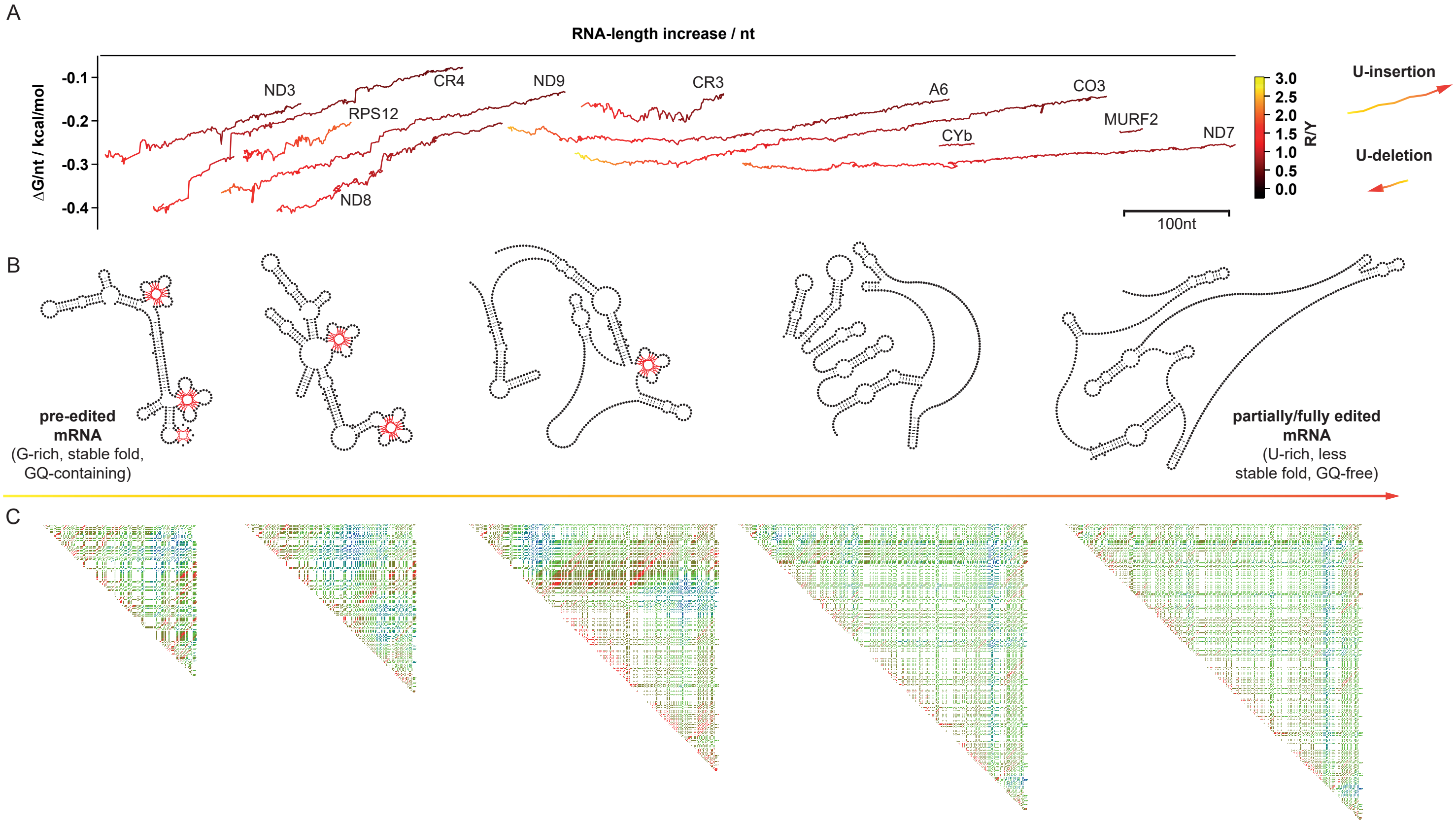


Supplementary Figure 1

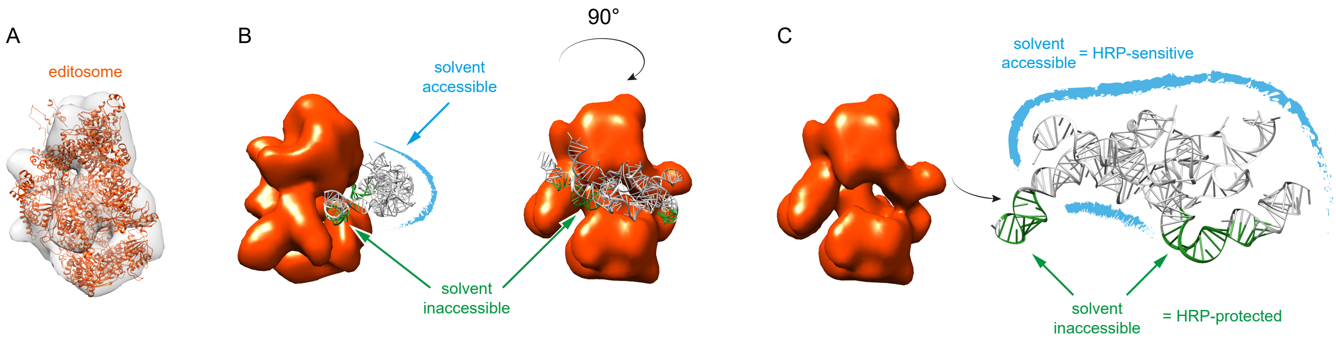


Supplementary Figure 2

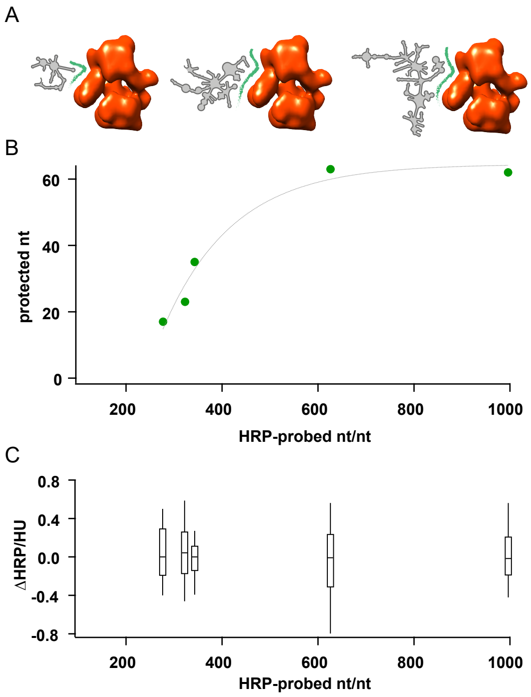


Supplementary Figure 3

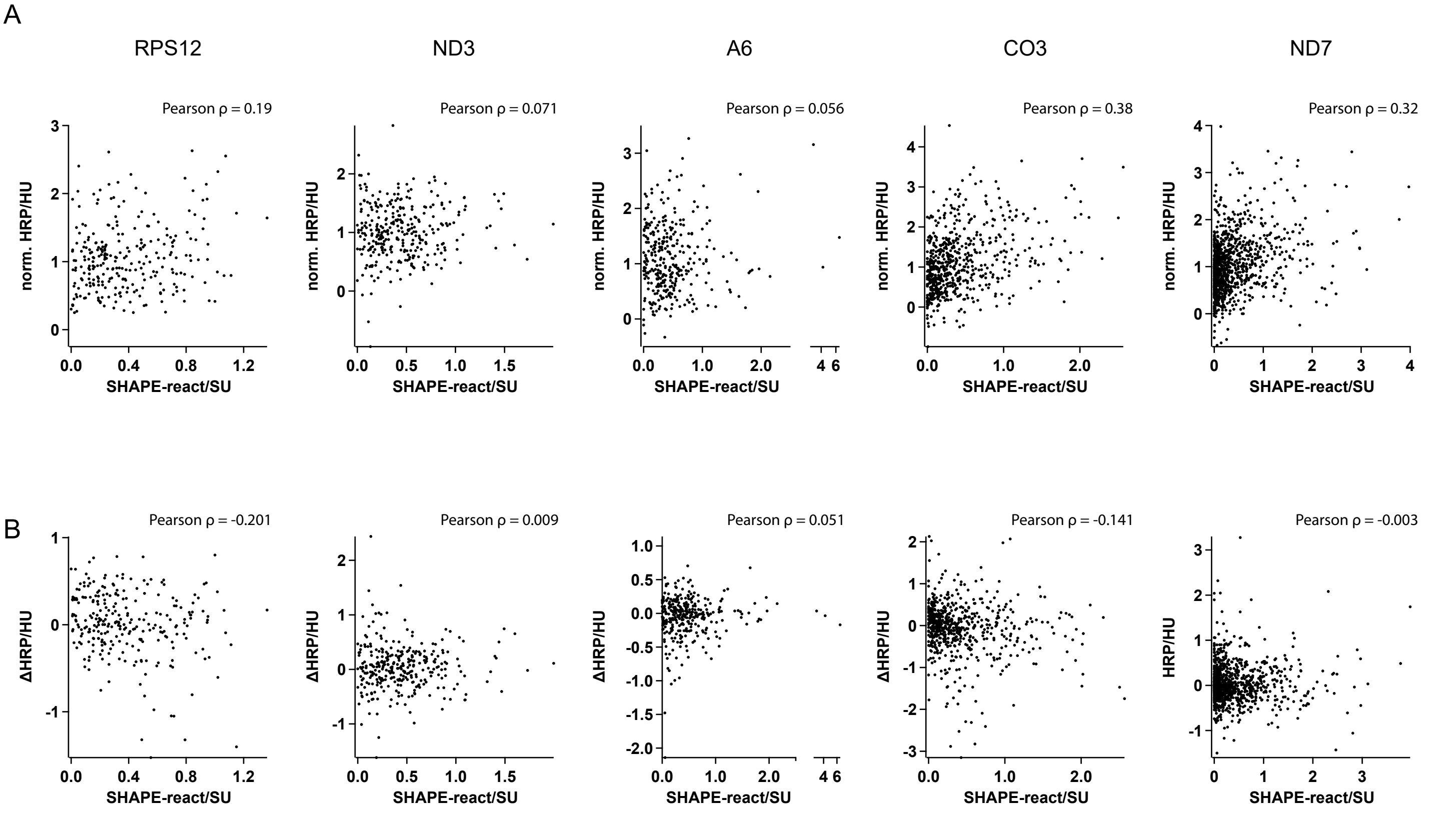


Supplementary Figure 4

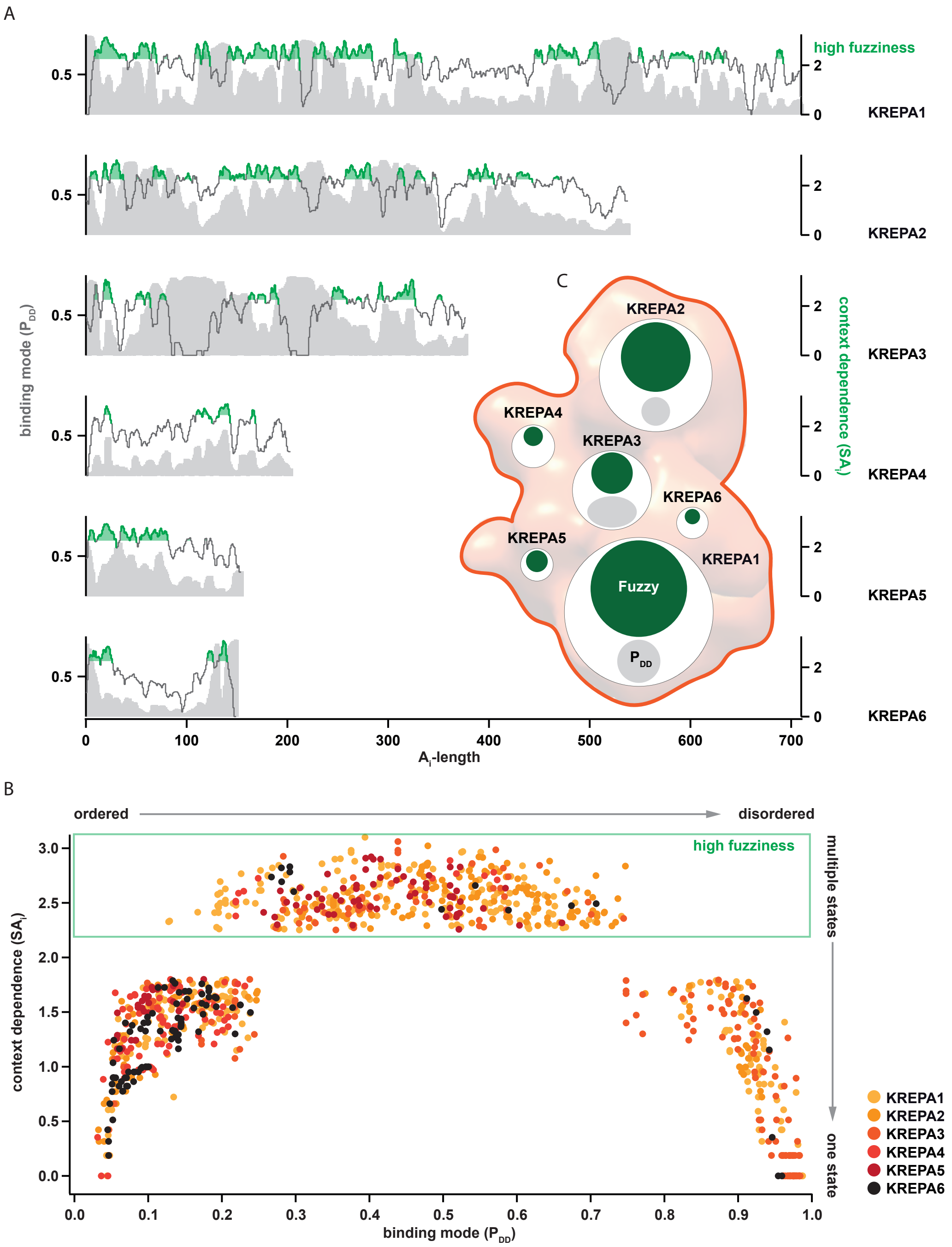


Supplementary Figure 5
